# String/Cdc25 phosphatase is a suppressor of Tau-associated neurodegeneration

**DOI:** 10.1101/2022.06.14.496126

**Authors:** Andreia C. Oliveira, Madalena Santos, Mafalda Pinho, Carla S. Lopes

## Abstract

Tau pathology is defined by the intracellular accumulation of abnormally phosphorylated Tau and is prevalent in several neurodegenerative disorders. The identification of modulators of Tau abnormal phosphorylation and aggregation is key to understand disease progression and develop targeted therapeutic approaches. In this study we identify String/Cdc25 phosphatase as a suppressor of Tau abnormal phosphorylation and associated toxicity. Using a *Drosophila* model of tauopathy we show that Tau dephosphorylation by Stg/Cdc25 correlates with reduced Tau oligomerization, brain vacuolization and locomotor deficits in flies. Moreover, using a disease mimetic model, we provide evidence that Stg/Cdc25 reduces Tau phosphorylation levels independently of Tau aggregation status and delays neurodegeneration progression in the fly. These findings uncover a role for Stg/Cdc25 phosphatases as regulators of Tau biology, that extends beyond their well-characterized function as cell-cycle regulators during cell proliferation, and point-out Stg/Cdc25 based approaches as promising entry points to target abnormal Tau phosphorylation.

## Introduction

Tauopathies are a group of neurodegenerative disorders defined by the intracellular accumulation and aggregation of Tau, a microtubule binding protein. These disorders include Alzheimer’s disease (AD), Parkinson, progressive supranuclear palsy and frontotemporal dementia (FTDP-17), among others (reviewed in Chung et al., 2021). Although the mechanisms leading to Tau aggregation and toxicity are still not fully understood, it is recognized that changes in protein structure/solubility lead to dimer and oligomer formation, which assemble into fibrillary structures, and large insoluble aggregates (Sivanantharajah et al., 2019).

Mutations in MAPT, the Tau encoding gene, and post-translational modifications (PTM), including phosphorylation, acetylation and ubiquitination, are the most common alterations associated with Tau pathology (Limorenko and Lashuel, 2022a; Tapia-Rojas et al., 2019). Recently, cryo-EM studies showed that cross-talk between PTMs underlies Tau structural diversity, impacts on fibril structure and correlates with distinctive tauopathies (Arakhamia et al., 2020; Fitzpatrick et al., 2017). Thus, the identification of Tau cellular partners that elicit changes in Tau structure and biology is key to understand pathophysiological mechanisms and prompt the development of effective therapeutic approaches.

*Drosophila* tauopathy models have been successfully used to uncover Tau interactors and investigate the molecular basis of Tau pathogenesis (Limorenko and Lashuel, 2022b). Pan-neuronal expression of wild-type or mutated human Tau (hTau) isoforms in flies recapitulates key pathological features of human tauopathies, including accumulation of abnormally phosphorylated forms of Tau, neuronal loss, progressive motor deficits and neurodegeneration. When expressed in the developing fly retina, hTau induces alterations in the external eye structure, characterized by the appearance of a rough eye phenotype that correlates with photoreceptor axons degeneration and loss of retinal cells (Jackson et al., 2002; Prüßing et al., 2013). This phenotype has been widely used in genetic screens and enabled the identification of cellular processes involved in Tau toxicity, which include Tau phosphorylation and proteolysis, cytoskeleton reorganization, chromatin regulation, and apoptosis (Feuillette et al., 2020; Hannan et al., 2016).

Unbalanced activity of kinases and phosphatases has been associated with Tau pathology. Interestingly, Shulman and Feany (2003) reported a genetic interaction between Tau and String, the fly Cdc25 homologue, when screening for suppressors of the rough eye phenotype induced by TauV337M, a mutation associated with frontotemporal dementia and Parkinsonism linked to chromosome 17 (FTDP-17) (Shulman and Feany, 2003). Phosphatases from the Cdc25 family (Cdc25A, B and C) are dual-specificity phosphatases whose function is associated with proliferating tissues, as key regulators of Cyclin-dependent Kinases (CDK) activity during cell division (Rudolph, 2007). Despite neurons in the normal adult brain have exited the cell cycle, Cdc25A, Cdc25B and Cdc25C phosphatases are expressed in this tissue, and display basal enzymatic activities (Ding et al., 2000; Vincent et al., 2001). Interestingly, increased expression and activity of Cdc25A has been reported in brain tissue from Alzheimer’s disease patients (Ding et al., 2000; Vincent et al., 2001). The role of CDC25 phosphatases in neurons is still unclear, as is their link to Tau biology and neurotoxicity.

We previously showed that String (Stg) phosphatase is expressed in photoreceptor neurons (Lopes and Casares, 2015). Here we use a fly tauopathy model to investigate the link between Stg/Cdc25 phosphatase and Tau and explore the neuronal function of Cdc25 phosphatases. We show that Stg/Cdc25 suppresses Tau-induced phenotypes, confirming the genetic interaction previously reported. This occurs with concurrent reduction of Tau phosphorylation levels revealing Stg/Cdc25 phosphatases as novel modulators of Tau toxicity *in vivo*.

## Results

### Stg phosphatase activity suppresses Tau-induced phenotypes

To explore the genetic interaction between Stg and Tau, we used the well-established fly tauopathy model *GMR-hTau2N4R* (hereafter *GMR-hTau*), which is based on the expression of the longest hTau isoform (2N4R) under the control of the GMR promoter. Accordingly, hTau is expressed in all cells of the developing retina, including photoreceptor neurons under GMR control (Ellis et al., 1993). To validate Stg as a neuron-specific modifier of hTau toxicity we used a *GMR-hTau* fly line that contains *elav*-Gal4 in the genetic background (*elav-Gal4; GMR-hTau*) and limits the expression of UAS-sequences exclusively in photoreceptor cells.

We used the external morphology of the retina as a readout of the Stg - hTau interaction (Fig 1 A - G). As previously reported (Jackson et al., 2002), when compared to control flies (*elav/+;* Fig. 1A), hTau expressing flies (*elav;GMR-htau/+; Fig. 1B)* display a rough eye phenotype with disordered ommatidia and missing or irregular mechanosensory bristles (Fig. 1B, Fig. S1). In addition, the retina is smaller as shown by morphometric analysis of circularity and length of the anterior-posterior (A-P) axis (Fig. 1 F, G). Stg expression in hTau expressing flies (*elav;GMR-htau/Stg*), the retina organization and size are restored (Fig. 1C, Fig. S1), with A-P length and circularity undistinguishable from control flies (Fig. 1 F, G). In contrast, knocking down Stg using RNA interference in hTau-expressing flies (*elav;GMR-hTau/Stg* ^*RNAi*^) has a minor impact on retina size and morphology of hTau expressing flies (Fig. 1 D, F, G).

**Fig 1.**
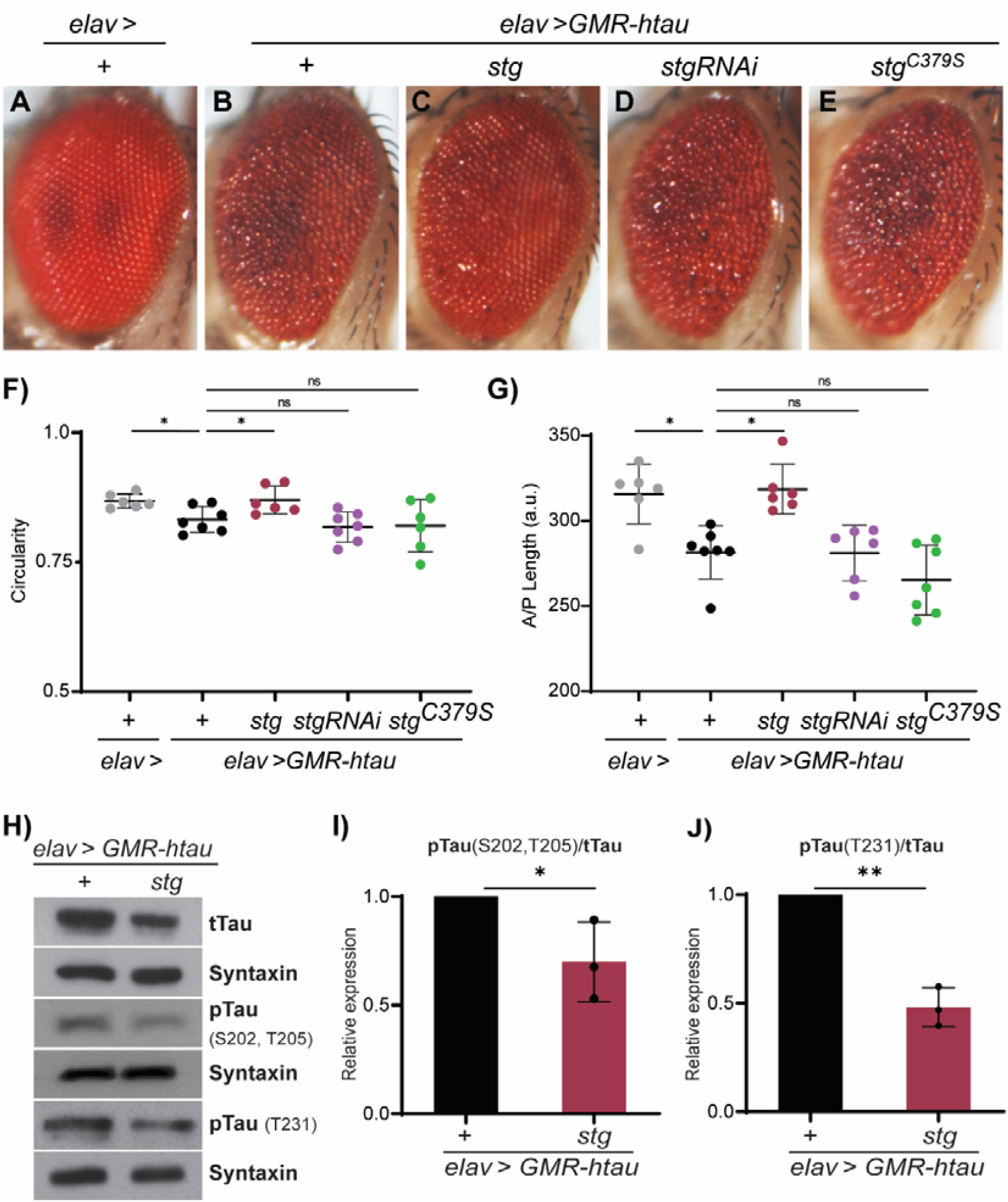
Stg phosphatase suppresses the rough eye phenotype and hyperphosphorylation associated with hTau expression in the fly retina. (A-E) Representative images of adult retinas (A) carrying elav-Gal4 or expressing (B) one copy of hTau (*GMR-hTau*), (C) hTau and Stg (*elav> GMR-hTau; stg*) (D) Stg^RANi^ (*elav> GMR-hTau; stg*^RANi^) or (E) a phosphatase dead Stg allele (*elav> GMR-hTau; stg*^*C379S*^). (F) Quantification of circularity of the retina of flies for the indicated genotypes. (G) Quantification of the anterior -posterior axis length for the indicated genotypes. Image quantification was performed on Fiji-ImageJ and the results were analyzed using one-way ANOVA with multiple comparisons; n≥6. (H) Western blot on protein extracts from fly heads of the indicated genotypes, blotted with total Tau and Tau phosphorylated at S202/T205 and T231. Syntaxin was used as loading control. (I, J) Quantification of the levels of phosphorylated Tau at S202/T205 (I) and T231 (J) normalized to total hTau levels. Data refers to three biological replicates. Data were analyzed using unpaired t-test with Welch’s correction. Error bars denote SD; n.s. indicates non-significant; * p < 0.05; ** p < 0.01; ****p < 0.0001.

Next, we asked if Stg phosphatase activity was required for the suppression of the *hTau* rough eye phenotype. The active site of Cdc25 phosphatases is highly conserved, and modification of the conserved cysteine in the active HCX_5_R site, was shown to abolish phosphatase activity in different models (Dunphy and Kumagai, 1991; Gautier et al., 1991; Rudolph, 2007; Sohn et al., 2004). Accordingly, we generated a Stg allele harboring a cysteine to serine (C379S) modification on Stg conserved active site (UAS-stgC379S; Stg phosphatase dead). Our results show that hTau; StgC379S flies show severe ommatidia disorganization, without significant changes in retina size compared to hTau expressing flies (Fig. 1E, F, G). Altogether, our findings confirm the Stg-hTau genetic interaction and show that Stg phosphatase activity is required to suppress Tau-associated phenotypes, leading us to propose that Stg/Cdc25 phosphatases may act as regulators of Tau toxicity *in vivo*.

### Stg activity reduces Tau phosphorylation levels

We next investigated if Stg impacts on Tau phosphorylation. Our analysis focused on phosphorylation of Ser202/Thr205 and Thr231 residues, which correlate with increased hTau aggregation propensity, AD clinical progression, and are used for diagnosis (Bibow et al., 2011; Hanger and Noble, 2011; Jeganathan et al., 2008; Wegmann et al., 2021). Accordingly, we compared the levels of total and site-specific phosphorylated hTau in hTau and hTau; Stg expressing flies (Fig 1. H-J). Western blot analysis shows that hTau phosphorylation levels at Ser202/Thr205 and Thr231 residues, are reduced by the co-expression of Stg in fly tissues (Fig 1 H). Quantification of the ratio between phosphorylated and total hTau indicates a reduction in phosphorylation levels in all residues under study (Fig.1 I, J). These results clearly show that Stg is able to counteract hTau phosphorylation in vivo.

### Endogenous Stg clusters with hTau in neurons

The reduced hTau phosphorylation levels detected upon Stg expression can be explained by direct Stg-hTau interaction. Alternatively, Stg can regulate hTau phosphorylation status indirectly, through regulation of Tau kinases and phosphatases. Since the association of protein phosphatases like Stg/Cdc25 with their substrates is transient and difficult to detect (Fahs et al., 2016; Sohn et al., 2004), we used in situ proximity ligation assay (*in situ* PLA) to probe Stg-hTau interaction. We used a stg allele expressing Stg - GFP fusion under stg endogenous promoter (*Stg-GFP, protein trap*) to evaluate Stg-Tau proximity in photoreceptors, avoiding false-positive interactions due to Stg overexpression. In third instar eye imaginal discs Stg-GFP is expressed at high levels in the proliferative/anterior domain and immediately posterior to the morphogenetic furrow (Fig. 2A). Immunolocalization studies in *elav;GMR-hTau/Stg-GFP* larvae show that hTau is expressed in cells posterior to the morphogenetic furrow, following the expression pattern of GMR promoter and overlapping Stg-GFP expression domain (Fig. 2B). PLA assays reveal the presence of *in situ* PLA signals in the domain co-expressing Stg and hTau (Fig. 2C), which indicates that Stg-GFP and hTau are found in close proximity in these cells. In contrast, no PLA signals are observed in eye-imaginal discs from *GMR-hTau; TM3,Ser* larvae confirming the specificity of in situ PLA-detection (Fig. S2).

**Fig 2.**
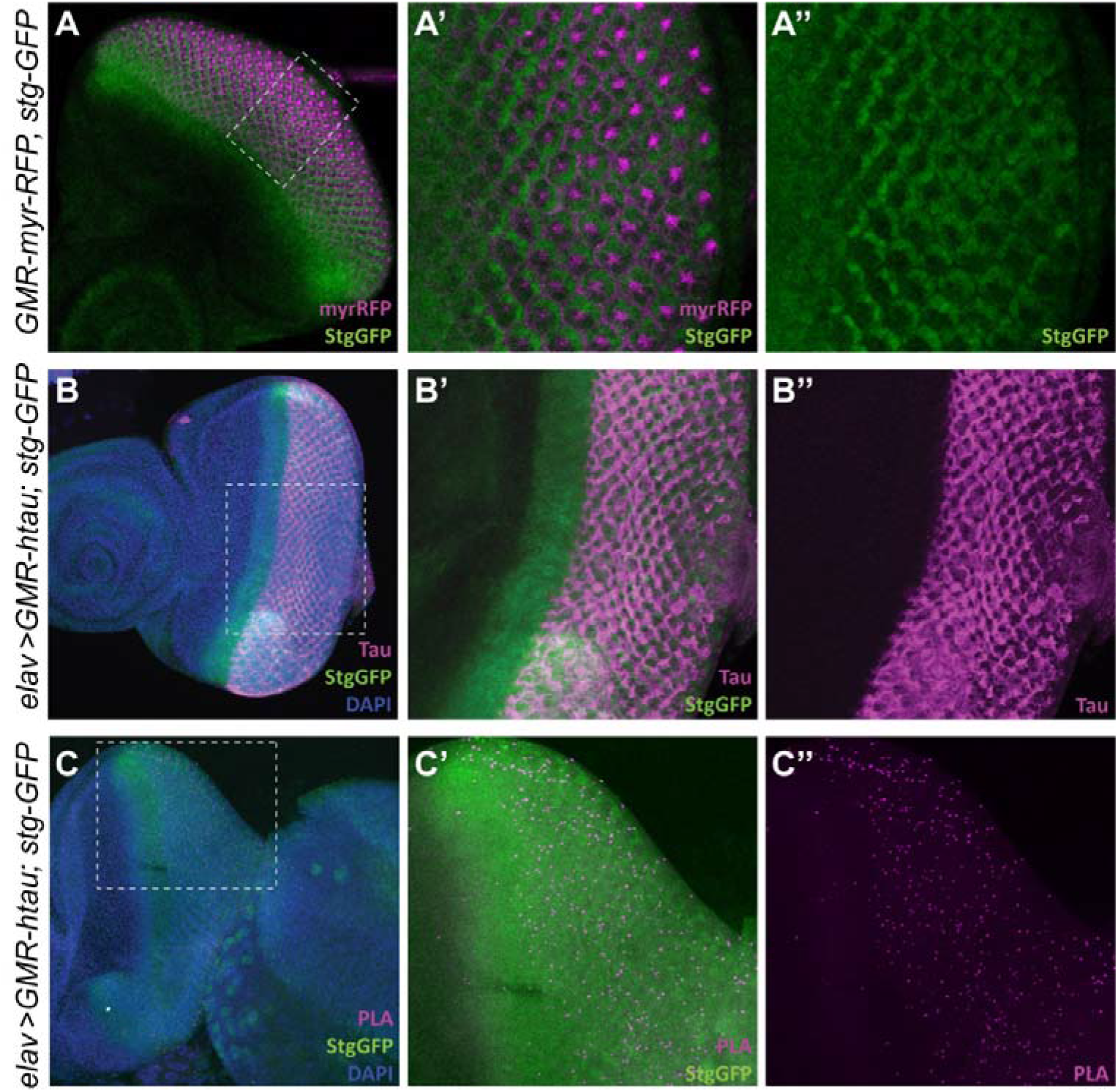
Endogenous Stg clusters with hTau. (A) Representative third instar eye imaginal disc from *GMR-myrRFP; stg-GFP* stained for GFP (green) to visualize Stg expression pattern. *GMR-myrRFP* (magenta) labels the GMR expression domain. (A’, A’’) Magnification of the region delimited by the dashed lines in A. (B) *GMR-hTau; stg-GFP* third instar eye imaginal disc, showing the expression pattern of hTau (magenta) and Stg-GFP (green). DNA (blue) was counterstained with DAPI. (B, B’’) Detail from the posterior domain of disc shown in B. (C) L3 eye imaginal disc from *elav>GMRhTau;stg-GFP* labeled for Stg (green) and showing PLA puncta (magenta), a readout of the interaction between endogenous Stg (green) and hTau. DAPI (blue) was used to counterstain DNA. (C, C’’) Magnification of the posterior region of the disc showing PLA puncta (magenta). In all images anterior is to the left.

### Stg suppresses Tau-driven neurodegeneration

Well-established neurodegeneration phenotypes like cell death (Jackson et al., 2002; Wittmann et al., 2001), brain vacuolization (Khurana et al., 2006; Mershin et al., 2004; Wittmann et al., 2001) and locomotor impairment (Sealey et al., 2017; Ubhi et al., 2007) are strongly associated with pan-neural expression of hTau and phosphorylation status (Nishimura et al., 2004). Therefore, we asked if Stg expression would modify neurodegeneration signatures in hTau-expressing flies.

To assay locomotor function we performed climbing assays in 8, 15- and 22-day-old flies using a countercurrent apparatus (Fig. S3). In this assay we evaluate the fly ability to climb-up a tube, in a set period of time, in 5 consecutive attempts (Inagaki et al., 2010). Flies that successfully climb-up the tube are transferred to a new tube and are challenged again. Thus, flies reaching the last tubes, 5 and 6 (group III), have increased climbing ability compared to those retained in tubes 1 and 2 (group I), despite given the same five climbing attempts. We observed that the climbing performance of hTau-expressing flies is significantly impaired (Fig. 3A, B), in agreement with previous observations (Sealey et al., 2017). By day 8, only 3% of hTau expressing flies reach tubes 5 and 6 (group III), while 77% are retained in group I (tubes 1 and 2) (Fig 3A). In contrast, at the same timepoint, more than 10% of hTau; Stg expressing flies complete the assay (group III) (Fig. 3A). Their behavior is similar to those observed in control flies. Moreover, the climbing -up probability for each genotype, given by the partition coeficiency (Cf), shows that overtime hTau; Stg flies perform better than hTau flies (Fig 3B). The improved locomotor function associated with Stg expression is sustained in time and by day 22 hTau; Stg expressing flies remain indistinguishable from controls (Fig 3B, Fig. S3). Although the climbing performance decreased with time in all genotypes, most likely due to age-related deterioration, Stg co-expression increased Cf to values similar to control, in all timepoints analyzed (Fig. 3B).

**Fig 3.**
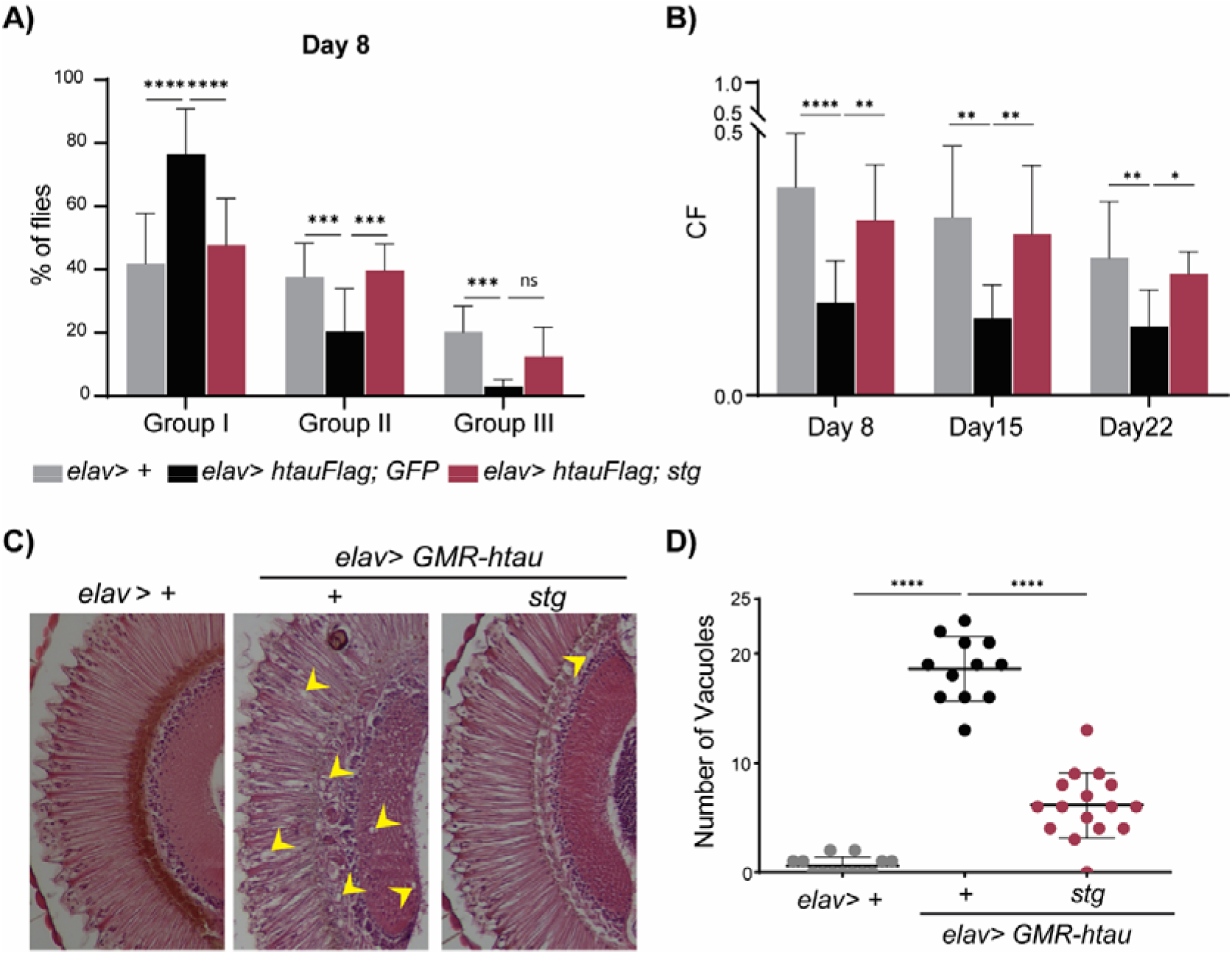
Stg rescues hTau-associated neurodegeneration phenotypes. (A) Quantification of the percentage of flies retained in group I (severe climbing impairment), II (middle climbing defects) and III (fittest flies) at day 8, for the genotypes indicated. Statistical significance was calculated using two-way ANOVA, multiple comparisons. (B) Graphic representation of the climbing index for 8, 15 and 22 for the indicated genotypes. Statistical significance was calculated using one-way ANOVA, multiple comparisons. (C) Representative cross-sections of adult retinas from control (*elav/+), hTau (elav> GMRhtau/+*) and hTau;Stg (*elav>GMRhtau;stg*) expressing flies, stained with hematoxylin and eosin. Yellow arrowheads indicate vacuoles. (D) Quantification of the number of vacuoles in the lamina with a diameter equal or superior to 3,5 μm for the genotypes indicated. Statistical significance was calculated using one-way ANOVA with Tukey’s multiple comparison test; n≥12. Error bars denote SD; * p <0.05; ** p <0.01; *** p <0.001; **** p <0.0001.

Next, we evaluated the internal morphology of the retina using standard hematoxylin-eosin staining (Fig. 3C). Analysis of adult head sections from GMR-hTau flies reveals severe disruption of the internal structure of the visual system, with strong vacuolization in both retina and lamina regions (Fig. 3C). Importantly, expression of Stg in photoreceptors improves significantly the internal structure of the retina of GMR-htau flies (Fig. 3C). In addition, quantification of the number of vacuoles between GMR-htau and GMR-htau; Stg expressing flies shows a strong reduction in vacuole number in GMR-htau; Stg 10-day old flies (Fig. 3D). These observations indicate that Stg expression suppresses Tau-associated neurodegeneration phenotypes and suggest that the dephosphorylation mediated by Stg reduces hTau cellular toxicity.

### Oligomerization and Tau spreading is impaired by Stg activity

During the analysis of adult head sections, we consistently detected the presence of vacuoles in the central brain region of *GMR-hTau* flies (Fig. 4A). This was unexpected since hTau expression is limited to cells within the posterior domain of the eye imaginal disc (Fig. 2B) and is not expressed in brain neurons. The occurrence of vacuolization in the central brain area is suggestive of spreading of toxic forms of hTau from the retina to neighboring regions in the fly head. To address this hypothesis, we quantified the number of vacuoles in the central brain region and apoptotic cells using the cleaved Caspase 3 marker, in all genotypes under study (Fig. 4B). Our analysis shows that St<u>g</u> expression reduces vacuolization tendency in hTau expressing flies (Fig. 4 A, B). Quantification of cell death, after cleaved -Caspase 3 labeling, reveals increased apoptosis in *GMR-hTau* relative to control flies (Fig. 4 C, D). Importantly, the number of apoptotic cells is significantly reduced upon Stg expression (Fig. 4 C, D). These findings support the hypothesis that hTau toxicity extends beyond the retina and imply spreading of hTau toxic forms in *GMR-hTau* tauopathy model. Accordingly, we evaluated the presence of oligomers in extracts from hTau expressing flies, as these are strongly implicated in hTau toxicity and the most amenable for spreading (Mudher et al., 2017). Dot blot analysis with A11, an oligomer specific antibody (Abskharon et al., 2020), reveals strong reactivity with GMR-hTau extracts (Fig. 4 E). This contrasts with lower reactivity, in GMR-hTau; Stg extracts, for equivalent levels of total protein. Altogether these findings show that Stg impact on hTau phosphorylation correlates with reduced oligomerization potential and neurodegeneration.

**Fig 4.**
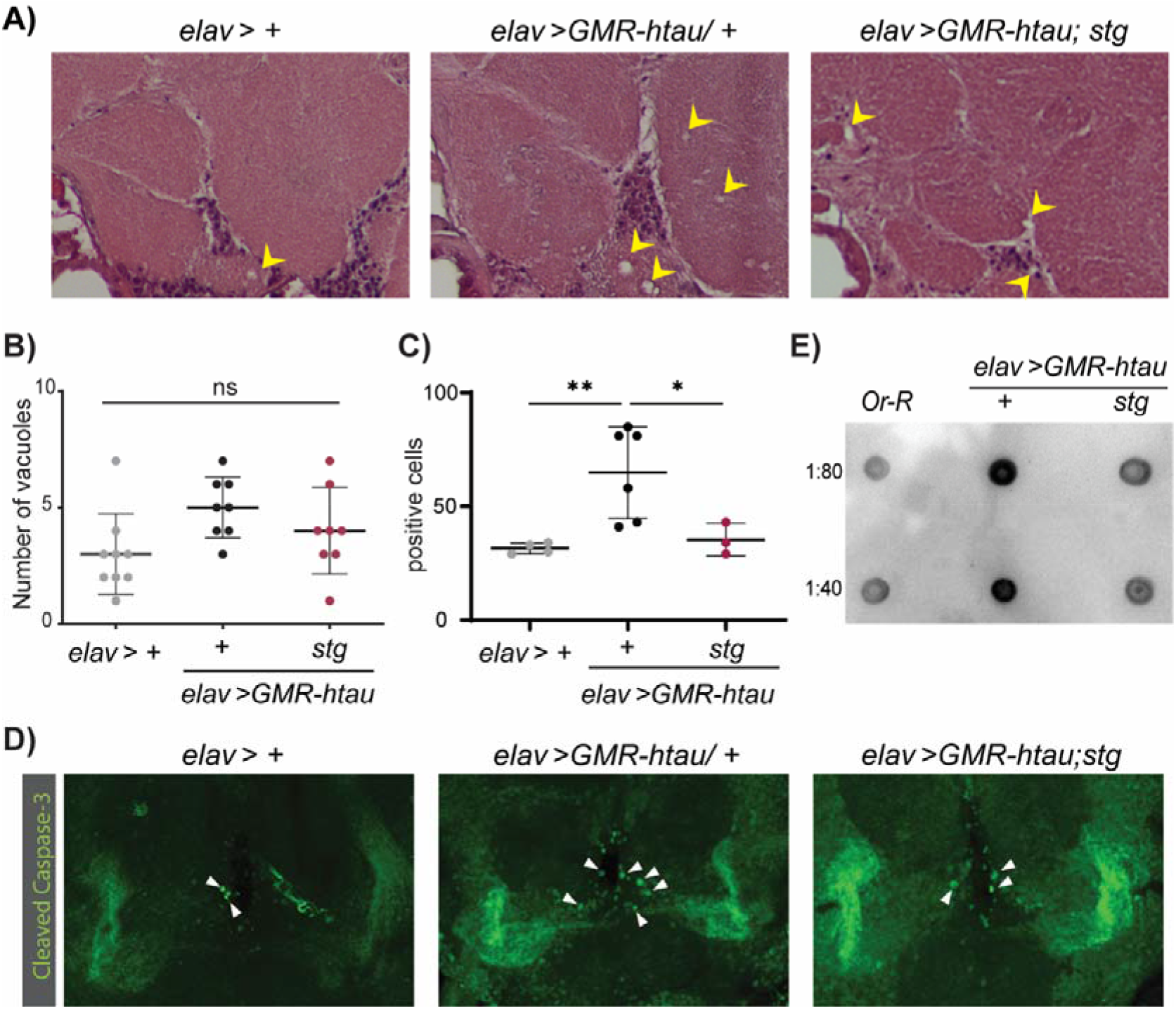
Stg reduces the levels of hTau toxic species. (A) Representative cross-sections of adult heads from control (*elav>+*), hTau (*elav> GMRhtau;+*) and hTau;Stg (*elav> GMRhtau,stg*) expressing flies, stained with hematoxylin and eosin. Vacuoles are indicated by yellow arrowheads. (B) Quantification of the number of vacuoles in the central brain region of 10-day old flies for the genotypes indicated; n≥ 8. (C) Quantification of Caspase-3 immunoreactive cells in brain of 10-day old flies; n≥3. (D) Representative images of adult fly brains stained for Caspase-3. Caspase-3 positive cells are indicated by white arrowheads. (E) Dot-blot analysis of oligomeric species in soluble fraction of extracts from fly heads for the indicated genotypes. Two dilutions of extract were analyzed. Statistical significance was calculated using one-way ANOVA. Error bars denote SD; n.s. indicates non-significant; * p <0.05; ** p <0.01.

### Stg suppresses neurotoxicity after Tau aggregation is established

The results presented thus far highlight Stg phosphatase as regulator of hTau phosphorylation and toxicity with effective physiological improvement in neurodegeneration progression. We next asked whether the up-regulation of Stg activity would be an advantage in a disease context, as this will allow the community to put forward therapeutic approaches built upon Tau regulation by Stg/Cdc25 phosphatases. To test this possibility, we assayed if overexpressing Stg after Tau hyperphosphorylation and aggregation are established would reduce Tau – associated toxicity. We used the Gal80^ts^ system to block the expression of UAS-Stg until *elav>GMR-hTau* individuals reach adult stage. Analysis of protein levels in head extracts from 10-day old flies, shows a significant reduction of hTau phosphorylated at Ser 202/Thr205 after Stg up-regulation (Fig. 5A, B). This clearly shows that Stg phosphatase can dephosphorylate hyperphosphorylated oligomeric forms of hTau *in vivo*.

**Fig 5.**
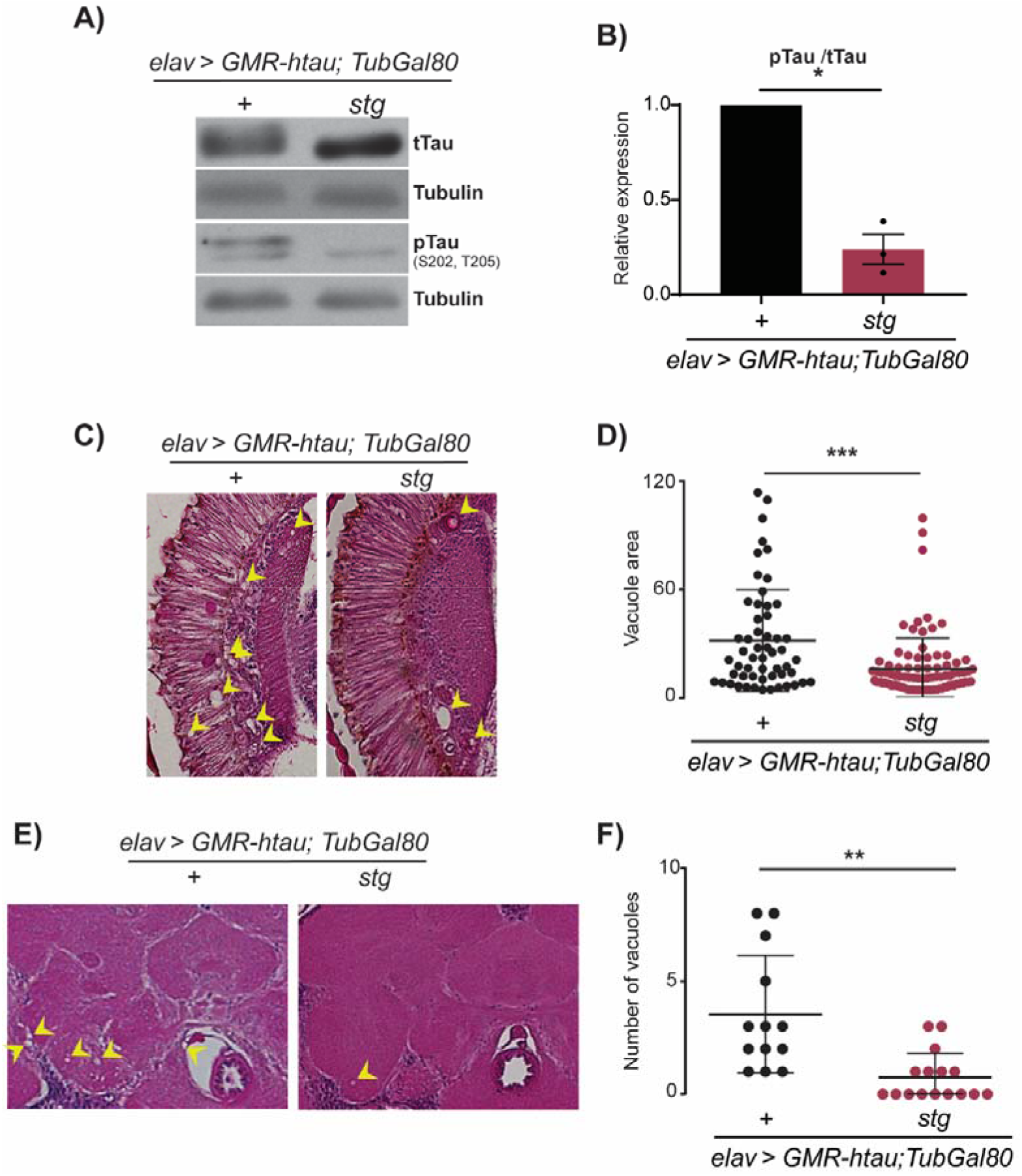
Stg promotes dephosphorylation of hTau toxic species delaying neurodegeneration progression. (A) Representative immunoblots of extracts from 10-day-old adult flies expressing hTau (*elav> GMR-hTau;TubGal80*^*ts*^) or hTau; Stg (*elav; GMR-hTau/stg;TubGal80*^*ts*^), probed with total and phosphorylated Tau. (B) Quantification of the ratio between Tau phosphorylated at Ser202/Thr205 and total hTau levels. Tubulin was used as loading control. Data represents three biological replicates. (C, E) Representative images from (C) retina and (E) central brain of 10 days old flies, of the indicated genotypes, stained with hematoxylin and eosin. Vacuoles are indicated by yellow arrowheads. (D) Quantification of vacuole size (diameter ≥ 3,5μm) in the retina and lamina regions. (F) Quantification of the number of vacuoles in the central brain region for the genotypes indicated. The results were analyzed using unpaired t-test with Welch’s correction, n ≥13. Error bars denote SD; * p < 0.05; ** p < 0.01; *** p < 0.0001.

We next asked if the lower levels of phosphorylated hTau correlate with slower disease progression, using vacuolization of the retina and brain as a readout (Fig. 5 C -F). As retina is completely formed when flies eclode, Stg overexpression failed to restore the internal structure of the retina in 10-day old flies (Fig. 5 C). However, it was sufficient to promote a reduction in vacuole area (Fig. 5 D). Analysis of the brain region indicates that Stg expression significantly reduces the number of vacuoles detected (from 3,5 to 0,75 vacuoles, on average) (Fig. 5 E, F). Overall, our findings show that Stg is able to promote dephosphorylation of aggregated and abnormally phosphorylated hTau.

## Discussion

Here we show that Stg, a conserved member of the Cdc25 phosphatases family is a suppressor of Tau-associated toxicity and neurodegeneration. Using a fly model of Tau2N4R tauopathy, we demonstrate that neuron-specific up-regulation of Stg activity reduces hTau phosphorylation levels and suppresses hTau-associated phenotypes, including rough eye, locomotor deficits and brain neurodegeneration. Our work provides evidence for an unambiguous Stg - Tau genetic interaction, expanding the former identification of Stg as a genetic modifier of TauV337M phenotype, a mutation associated with familial frontotemporal dementia (Shulman and Feany, 2003). Importantly, these findings highlight a novel role for Stg/Cdc25 phosphatases as Tau regulators, with impact on disease outcome. Cdc25 phosphatases (Cdc25A-B, and C) play essential roles during normal cell division as regulators of CDK activity, and their increased activity is associated with cell proliferation and tumor progression (Boutros et al., 2006; Boutros et al., 2007). Interestingly, increased expression of Cdc25A and B was detected in samples from AD disease brains, and both were shown to accumulate in the cytoplasm of degenerating neurons (Ding et al., 2000; Vincent et al., 2001). Cdc25 up-regulation correlated with increased phosphatase activity towards Cdk1, and the occurrence of mitotic figures in degenerating neurons (Ding et al., 2000; Vincent et al., 2001). Likewise, increased activity of Cdc25A was linked to hypoxia-induced neuronal death, and correlated with increased pRB phosphorylation (Iyirhiar et al., 2017). The up-regulation of cell cycle genes, including Cdc25, is regarded as a neuronal response to insults, as DNA damage, that promotes aberrant cell-cycle re-entry, and ultimately leads to neuronal death (Chang et al., 2012; Marlier et al., 2020). However, our findings suggest that increased expression of Cdc25 in neurons can be neuroprotective. In agreement, Cdc25 overexpression promotes regeneration in *Drosophila* sensory neurons after physical injury, whereas Cdc25 knockdown impedes regeneration (Li et al., 2021). In fact, Cdc25 participates in the evolutionary conserved Piezo-Atr-Chek1-Cdc25 inhibitory regeneration pathway. We propose that Cdc25 may play a neuroprotective role during early steps of Tau-induced neurodegeneration, in response to compromised axonal dynamics and homeostasis, while at later stages of disease the co-occurrence of DNA damage elicits cell-cycle re-entry culminating in neuronal death.

Consistent with a potential homeostatic function for Cdc25 phosphatases in neurons, Cdc25A and B expression has been reported in human brain samples (Ding et al., 2000; Vincent et al., 2001), mouse (Iyirhiar et al., 2017) and rat (Chang et al., 2012). The Stg/Cdc25-Tau genetic interaction, herein described, supports a function as cytoskeleton regulators. Importantly, the functional conservation between Stg and mammalian Cdc25 phosphatases predicts that Stg-Tau interaction is likely conserved throughout evolution. Accordingly, we propose that Stg/Cdc25 comprise novel Tau phosphatases with strong impact on Tau biology and pathology, whose activity deserves further studies. Thus, Stg/Cdc25 adds to the list of key Tau regulators like Shaggy/GSK3-B (Jackson et al., 2002), Par-1/MARK (Ambegaokar and Jackson, 2011; Nishimura et al., 2004), MAPK and PP2A phosphatases (Shulman and Feany, 2003) identified in flies (reviewed in (Gistelinck et al., 2012; Hannan et al., 2016; Limorenko and Lashuel, 2022b).

Our findings show that the Stg/Cdc25 phosphatase counteracts hTau-associated toxicity. We detected a correlation between reduction of Tau phosphorylation levels mediated by Stg and suppression of neurodegeneration phenotypes. In addition, Tau dephosphorylation correlated also with reduced oligomerization in fly brains. These findings suggest that Stg/Cdc25 promotes dephosphorylation of residues that impact directly on Tau structure and biology. Accordingly, we detect reduced phosphorylation levels at hTau residues abnormally phosphorylated in AD brains (Hanger and Noble, 2011). The close proximity detected among endogenous Stg and hTau, alludes a direct interaction between the two proteins. Therefore, we put forward the hypothesis that Tau dephosphorylation is likely the outcome of direct Stg/Cdc25 activity. However, indirect regulation of Tau phosphorylation status, through modulation of Tau kinase(s) activity cannot be ruled out.

Importantly, our observations unveil the potential of therapeutic approaches based on Cdc25-Tau interaction. In contrast to studies based on co-expression approaches, during fly development or restricted to adult stages (Cowan et al., 2010; Fernandez-Funez et al., 2015; Hannan et al., 2016), here we use a paradigm of disease and induce Stg/Cdc25 expression after high levels of phosphorylated Tau build-up in neurons. This approach enabled us to show that Stg/Cdc25 is able to promote dephosphorylation of highly-phosphorylated toxic forms of hTau. Moreover, we observed a reduction in vacuolization levels in the brain of 10-day old flies, in support of lagged neurodegeneration. This occurs albeit the maintenance of high levels of hTau protein. These findings show that it is the presence of highly phosphorylated soluble hTau that is detrimental to neurons, in agreement with other studies in flies and mice (Cowan et al., 2015; Feuillette et al., 2010).

Our results are in line with the hypothesis that hTau abnormal phosphorylation increases hTau toxicity and spreading potential. In flies only the lamina and medulla neuropils receive direct input from photoreceptor neurons (Fischbach and Dittrich, 1989; Sanes and Zipursky, 2010), however, we detected increased cell death and vacuolization in the central brain of *GMR-hTtau* flies. This was unexpected since in the GMR-hTau model, Tau expression is restricted to the presynaptic terminal of photoreceptors. Thus, brain vacuolization could only be explained by spreading of Tau-toxic forms, most likely oligomeric Tau, to other brain regions. Importantly, Tau dephosphorylation promoted by Stg correlated with reduced apoptosis and vacuolization levels in the brain of hTtau flies. Similar findings were reported in mice (Hu et al., 2016) were it was proposed that dephosphorylation of AD– hyperhosphorylated Tau shows reduced propagation in the brain. Although the cellular mechanism(s) underlying trans-cellular Tau propagation remain to be fully identified, *Drosophila* models as the one used in this study, can provide important contributions to the mechanisms (synaptic vesicles, endocytosis, diffusion of free Tau or oligomeric forms) and molecular players involved. We expect that future studies following-up on our findings will establish Cdc25 phosphatases as important regulators of Tau biology and conceivable venues to explore in the development of effective therapeutic approaches.

## Materials and Methods

### Fly strains and genetics

Flies were maintained in standard media, at 25ºC under a 12:12 light-dark cycle, unless indicated. The strains *elav* ^*C155*^ GAL4 (BDSC 458), *UAS-stg* (BDSC 4777), *UAS-stg*.*HA* (BDSC 56562), *elav*^*C155*^GAL4; GMR-hTau (BDSC 51360), tubGAl80^ts^ (BDSC 7017), *UAS-stgRNAi* (BDSC 34831), *Stg-GFP* (BDSC 50879), GMR::myrRFP (BDSC 7121), Oregon R were obtained from the Bloomington stock Centre (NIH P40OP018537). *UAS-hTau-Flag* was previously described (Kosmidis et al., 2010). Standard genetic techniques and fly lines carrying balancers on the second and third chromosomes were used to generate the different genetic backgrounds. Control flies in all experiments were as closely related to the experimental flies as possible. *elav*^*C155*^ Gal4 flies were crossed to *Oregon R* to generate heterozygous control alleles. Flies of genotypes containing the Gal4/UAS/Gal80^ts^ constructs were grown at 18°C, and transferred to 29°C as 1-3 day-old adults, to allow expression of the Gal4, and aged until use in histological or protein analysis. A list of the genotypes analyzed is provided in Supplementary Table 1. The phenotype of adult retinas was imaged in a stereomicroscope Stemi 2000 Zeiss equipped with a Nikon Digital SMZ 1500 camera, with a 50X magnification.

### Generation of transgenic flies

The *UAS-Stg*^*C379S*^ transgenic flies were generated by replacing the conserved cysteine within the catalytic domain to serine using the NZY Mutagenesis kit according to manufacturer’s protocol. The following oligonucleotides were used to amplify stg coding sequence from LD47579 plasmid, stgPPAseFW (5’-CAACATCATTATCTTCCACGCCGAATTCTCCTCGGAGCGT-3’) and stgPPAseRW (5’-ACGCTCCGAGGAGAATTCGGCGTGGAAGATAATGATGTTG-3’), followed by DpnI digestion of parental DNA (LD47579). After transformation of DH5α competent cells, plasmid DNA was amplified and at least three clones were selected for sequencing analysis. *EcoR1-XhoI* digestion was used to clone *stg*^*C379S*^ sequence into *pUAST-attB*. Transgenic flies were obtained after site-specific integration of *UAST-Stg*^*C379S*^ on the 3rd chromosome (3R-86F) (Bischof et al., 2007).

### Protein extraction and analysis

For protein analysis, adult flies of the appropriate genotypes were aged for 8-15 days, at appropriate temperature, flash frozen in liquid nitrogen and stored at -80ºC. Flies were decapitated by vigorously vortexing for 15 seconds, and heads were homogenized in RIPA buffer (50 mM Tris-Hcl, 150 mM NaCl, 1%Triton), supplemented with protease and phosphatase inhibitors (Roche) in dry ice. Extracts were incubated for 1h at 4ºC with rotation and centrifuged at 2000 g for 20 minutes, to separate soluble from insoluble fractions. Total protein content was quantified by the Lowry Method (DC™ Protein Assay, Bio-Rad, CA, USA). Fifteen micrograms of soluble extract were loaded in 10% SDS-polyacrylamide gels and transferred onto nitrocellulose membranes, for western blot analysis. Extracts were loaded in replicated gels and probed in parallel with total Tau and phosphorylation-specific Tau antibodies. Membranes were blocked during 1 hour with TBS 0,1% Tween containing either 5% low-fat milk or 5% BSA (Sigma). Primary antibodies were diluted in TBST 0,1%Tween with 3% blocking agent: phospho-Tau (Ser202, Thr205) (AT8) (1:2000, Termofisher Scientific), phospho-Tau (Thr231) (AT180) (1:2000, Termofisher Scientific), total Tau (5A6) (1:6000, DSHB), anti-Syntaxin (1:500, DSHB) and anti-Tubulin (1:10000, DSHB). Goat anti-rabbit and goat anti-mouse horseradish peroxidase-conjugated secondary antibodies (Amersham) were diluted at 1:10000 and 1:15000, respectively, in TBST 0,1% Tween containing 3% low-fat milk. Signal was detected using Clarity Western ECL Substrate reagent (Bio-Rad Laboratories) according to manufacturer’s instructions. A GS-800 calibrated densitometer with Quantity One 1-D Analysis Software 4.6 (Bio-Rad Laboratories) was used for quantitative analysis of protein levels. Western blots were repeated at least 3 times with biological replicates and representative blots are shown.

### Dot Blot

For dot bolt analysis, 10 fly heads were homogenized in RIPA buffer supplemented with protease and phosphatase inhibitors. After a 5 min centrifugation at 10000 g, samples were diluted in 1% Glycerol/PBS. 3μl of protein were dotted in nitrocellulose membrane. After blocking in 5% Milk /TBS-0,1%, the membrane was incubated with anti-amyloid oligomer A11 antibody (1:500, AB9234 Merck) overnight at 4°C, and handled for protein detection following the protocol described above. Images were acquired in ChemiDoc XRS (Bio-Rad).

### Histology analysis

Female flies were anesthetized on ice, immobilized in a histology collar and fixed with Carnoy’s fixative, overnight at 4ºC. Tissue was dehydrated in ethanol series prior to paraffin embedding and microtome sectioning as described in (Sunderhaus and Kretzschmar, 2016). Serial sections (5 μm) from the entire head were stained with hematoxylin and eosin and examined by bright-field microscopy. Images were captured with an Olympus DP72 microscope. Total vacuole number and vacuole area were quantified, in the central brain, retina and lamina, in at least 8 flies per genotype.

### Immunofluorescence analysis

Third instar larval tissues are dissected and fixed with 3,7% formaldehyde in PBS for 20 min, followed by washes in PBS 0.1% Triton. The primary antibodies used were: anti-GFP (A11122, 1:1000, Invitrogen) and anti-Tau (5A6, 1:5000, DSHB). The secondary antibodies were conjugated with Alexa Fluor dyes (ThermoFisher Scientific). Brains from 10 day -old flies were dissected in cold PBS and fixed in 3.7% formaldehyde in PBS for 30 min. Samples were washed three times in PBT 0.5% [1⍰× PBS⍰+⍰0.5% Triton X-100 (Sigma-Aldrich)] and incubated in blocking buffer (PBT 0.5%⍰+⍰0.5% BSA⍰+⍰0.5% FBS) for 1 h 30 min at room temperature. Incubation with primary antibodies anti-Cleaved caspase 3 (Asp 175) (1:50, Cell Signaling) and anti-ELAV (1:400, DSHB) was performed in blocking buffer overnight at 4 °C. Secondary antibodies conjugated with Alexa Fluor dyes (ThermoFisher Scientific) were diluted 1:500 in PBT 0.5% and incubated for 3 h at room temperature. Tissues were mounted in Fluoromount-G™ Mounting Medium (ThermoFisher Scientific). Samples were imaged on a Leica SP5 confocal microscope and images were processed with ImageJ (NIH).

### Proximity-ligation assay (PLA)

The Duolink In Situ Red system (92101, Merck) was used to detect Stg-Tau interaction in vivo, following the manufacturer’s protocol. Briefly, third instar larvae eye imaginal discs were dissected and fixed as described previously and incubated with Duolink Blocking Solution for 1 h at 37°C. Incubation with the primary antibodies anti-GFP (A11122, Invitrogen) and anti-hTau (5A6, 1:5000, DSHB) was performed ON at 4 C. Samples were washed twice with Duolink Wash Buffer A, and incubated with MINUS (anti-mouse) and PLUS (anti-rabbit) Duolink PLA Probes for 1 h at 37°C. Samples were Duolink Wash Buffer A prior to the ligation (1h at 37°C) and amplification (1h30h at 37°C) steps. Eye imaginal discs were mounted in Duolink In Situ Mounting Medium with DAPI (MERCK). Images were acquired using Leica SP5 confocal.

### Climbing assays

Climbing assays were performed in a counter-current apparatus equipped with six chambers, as described in Inagaki et al. 2009. Newly hatched flies of the appropriate genotypes were allowed to mate for 2 days, and after separated by gender in groups of 10-15 flies and aged at 25ºC. At each time point, flies were placed into the first chamber, tapped vigorously to the bottom and given 15 secs to climb upwards (approximately 10 cm), reaching the upper tube. The flies that successfully reached the upper tube were shifted to a new chamber, and both sets of flies were given another opportunity to climb upwards, in 15 secs. This procedure was repeated for a total of five times, and the number of flies in each chamber was counted. Flies were classified into three performance groups according to the locomotor ability displayed: flies that remained in tubes 1 and 2 were considered to have severe locomotor impairment (group I); those remaining in tubes 3 and 4 were considered to have moderate locomotor impairment (group II), while those that reached the last tubes, tubes 5 and 6, - group III - performed well in the climbing assay. Climbing assays were performed at days 8, 15 and 22, and partition coefficient (Cf) was calculated, according to (Inagaki et al., 2010). The partition coefficient (Cf) represents the probability of flies to climb up. At least 100 flies were used per genotype.

### Scanning Electron Microscopy

For SEM analysis, 1-day old flies were dehydrated through incubation in ethanol series (25%, 50%, 75%, 100%), incubated with hexamethildizilasane (HMDS, Sigma), air-dried, mounted in SEM stubs and coated with Au/Pd thin film, by sputtering using the SPI Module Sputter Coater equipment. Imaging was performed using a High resolution (Schottky) Environmental Scanning Electron Microscope with X-ray microanalysis and Electron Backscattered Diffraction analysis: Quanta 400 FEG ESEM/EDAX Genesis X4M. The images were acquired with 400X and 1000X magnification.

### Statistical Analysis

For western blot analysis, significance was determined using unpaired t-test with Welch’s correction, two-tailed. For climbing assay, the significance was measured using two-way ANOVA with Tukey’s multiple comparison test. For vacuole analysis, one-way ANOVA with Tukey’s multiple comparison test (Fig. 3,4) or unpaired t-test with Welch’s correction, two-tailed (Fig. 5) were used to determine significance. All statistical analyses were performed using GraphPad Prism, version 6.0 (GraphPad Software v8.4). All experiments were performed with biological triplicates. Significance is defined as p<0.05 and bars in graphs represent the mean ± SD.

## Acknowledgments

We thank F. Janody for support and critical comments on the manuscript, and members of the Janody and Relvas laboratories for important discussions. The Rat-Elav-7E8A10 anti-Elav developed by Rubin, G.M., mouse anti-Syntaxin 8C3 developed by Benzer S./ Colley N., were obtained from the Developmental Studies Hybridoma Bank, created by the NICHD of the NIH and maintained at the University of Iowa, Department of Biology, Iowa City, IA 52242. The authors acknowledge the support of the i3S Scientific Platform ALM scientific, member of the national infrastructure PPBI - Portuguese Platform of Bioimaging (PPBI-POCI-01-0145-FEDER-022122). This work was financed by FCT - Fundação para a Ciência e a Tecnologia (POCI-01-0145-FEDER-007274; NORTE-01-0145-FEDER-000008). ACO holds a PhD fellowship from FCT (SFRH/BD/130119/2017).

---

**Fig. S1.**
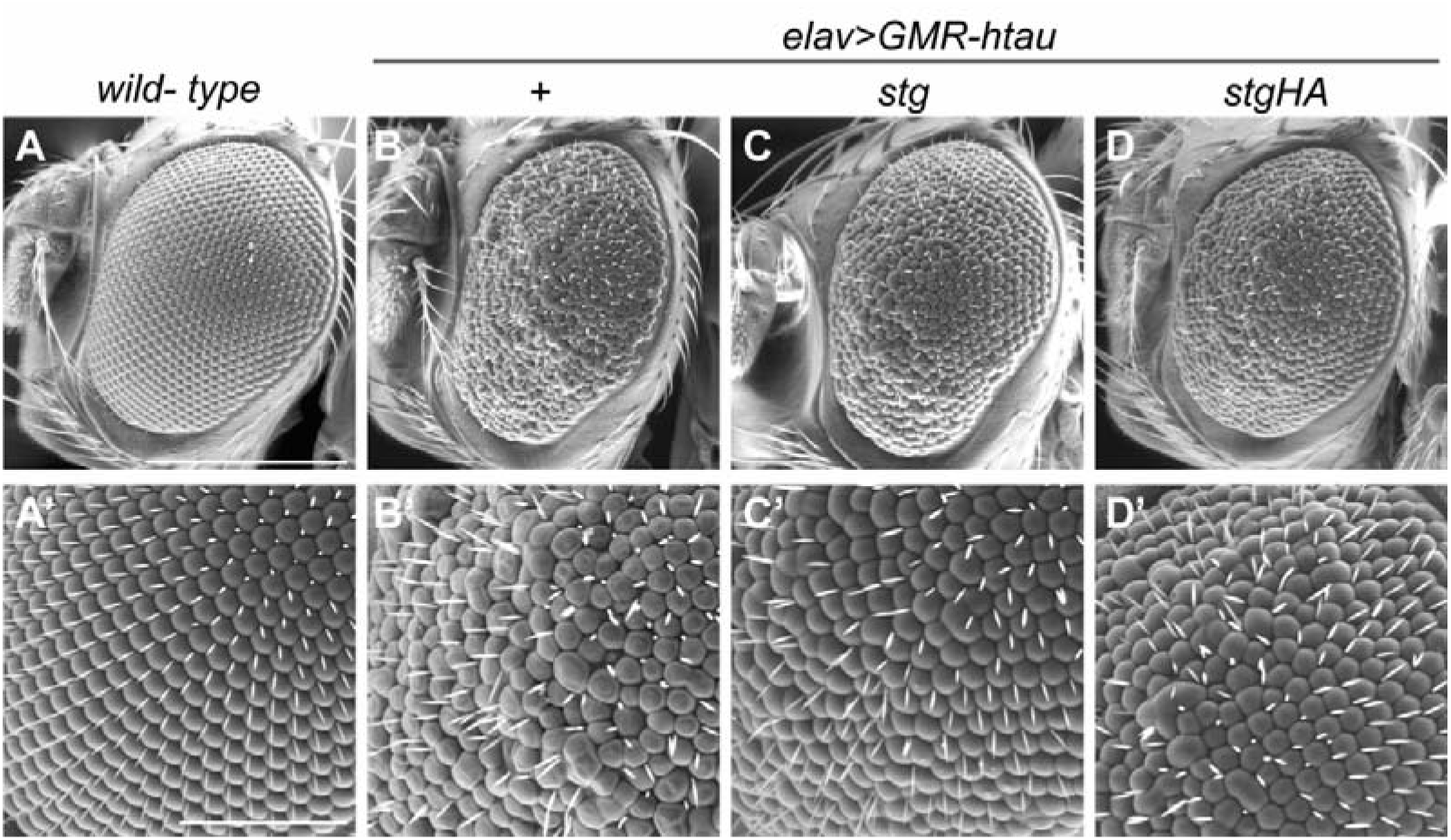
Stg suppresses the rough eye phenotype of hTau expressing flies. (A – D) Representative images of the retina of (A) control flies, (B) flies expressing one copy of hTau (*elav>GMR-hTau; +*), and (C,D) flies co-expressing Stg and hTau (*elav>GMR-hTau; stg, and elav>GMR-hTau; stg-HA)*. (A’-D’) magnified views of A, B, C and D, respectively.

**Fig. S2.**
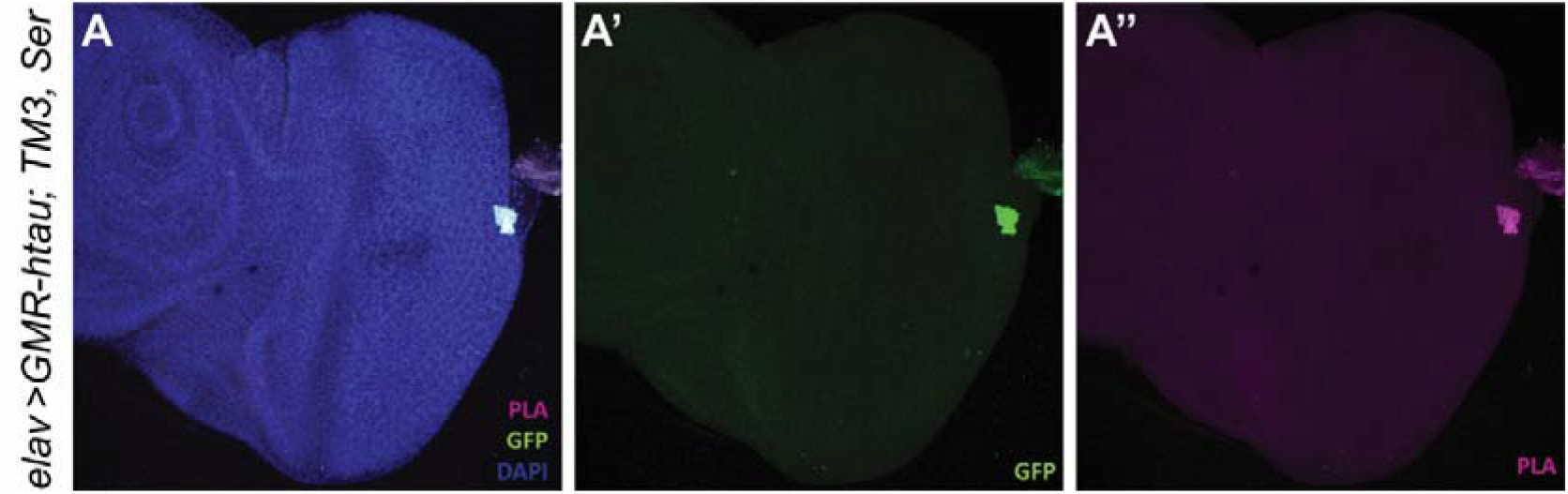
Negative control for PLA experiment. Representative images of eye imaginal discs from *elav> GMRhTau;TM3,Ser* larvae.

**Fig. S3.**
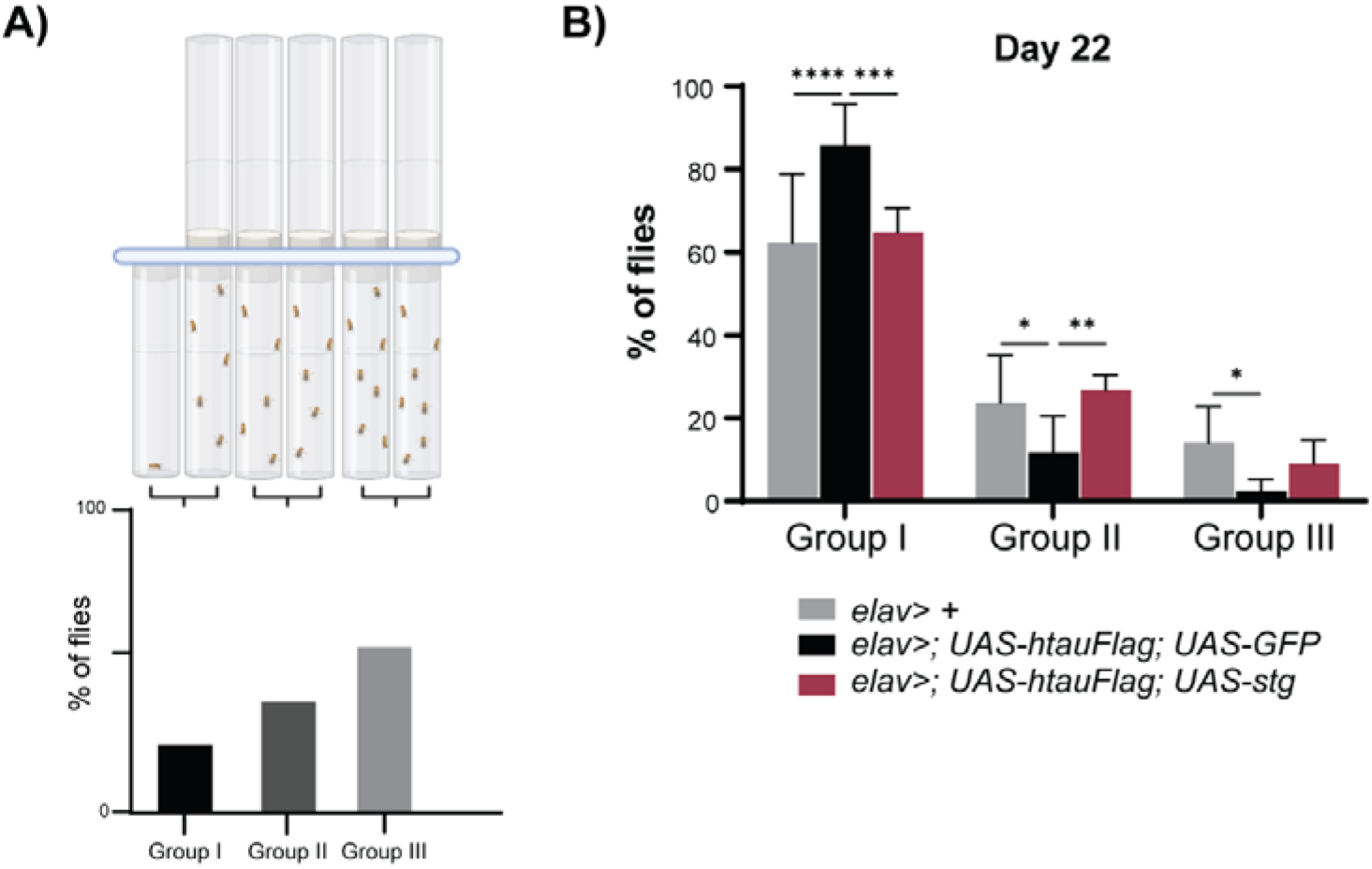
Graphical representation of the climbing assay with countercurrent apparatus. (A) Final distribution of the flies in the countercurrent apparatus represented as a bar graph. (B) Quantification of the percentage of flies in each group for the genotypes indicated, at day 22.

